# Quantification of *Trypanosoma Brucei* social motility indicates different colony growth phases

**DOI:** 10.1101/2024.05.03.592443

**Authors:** Andreas Kuhn, Timothy Krueger, Magdalena Schüttler, Markus Engstler, Sabine C. Fischer

## Abstract

*In vitro* colonies of the flagellated parasite *Trypanosoma brucei* grow in characteristic fingering instability patterns. The underlying cause of this this type of collective migration and the associated behavior called social motility remains a topic of intense debate in the scientific community. In order to study these mechanisms in detail, it is crucial to develop quantitative methods to assess and measure social motility patterns at the level of the whole colony beyond qualitative image comparisons. Here, we show a quantification of the growth process based on two scale free metrics designed to quantify the shape of two-dimensional colonies. While initially developed for yeast colonies, we adapted, modified and extended the analysis pipeline for the *Trypanosoma* system. Combining the quantitative measurements with colony growth simulations based on the Eden model, we discovered two distinct growth phases in social motility showing colonies: In the first phase, the colonies shown mainly circular expansion and later switch to an almost exclusive finger growing phase. These phases are robust when increasing the number of cells as well as upon partial inhibition of the finger formation. A newly developed anisotropy index shows that upon partial inhibition the anisotropy in the colony increases over time. Our results provide objective measurements that help advance the understanding of social motility of trypanosomes. Furthermore, our approach is suitable as a blueprint for investigations of other colony forming cells such as yeast or bacteria.

## 1. Introduction

Densely packed Trypanosma brucei colonies on agarose surfaces exhibit a phenomenon known as social motility (SoMo) [1]. These colonies initially manifest as circular structures but expand into characteristic flower-like patterns, featuring “petals” (often referred to as fingers or protrusions) displaying a distinct perpendicular alignment away from the colony’s center.

Numerous studies have delved into various facets of this phenomenon over the years. In 2014, Imhof et al. [2] demonstrated that SoMo is characteristic of the early procyclic form of *Trypanosoma brucei*. Subsequently, a pivotal role of intracellular cAMP levels in regulating SoMo [3, 4] has been proposed. Further, a correlation between the ability to collectively migrate *in vitro* and the development of trypanosomes in the tsetse fly have been established [5]. As a potential method of communication between cells, necessary for the proposed social aspect of the collective motion, secreted exosomes have been shown to have a repulsive effect on active colonies [6]. Finally, an actual positve chemotaxis was proposed to be caused by bacterial colonies, [7], which has later been attributed to pH taxis. [8]. The first quantification of single cell motility during collective migration, surprisingly showed minimal correlation. These results highlighted the importance of as yet unclear physical parameters underlying the phenomena of SoMo. [9].

Despite the noteworthy findings in these studies, a crucial aspect has been overlooked: a quantification of the investigated SoMo phenomenon at the level of the whole colony. Previous attempts have primarily relied on binary manual classification of SoMo-negative and -positive colonies. Various metrics have been explored, including cell number and density measurements [2, 5], finger or protrusion counts [3, 4, 5], and changes in distance between interacting colonies [6], alongside with a more sophisticated chemotactic index [7] and single-cell velocity measurements together with summary statistics [7, 9]. However, most of these quantifications only capture specific facets of the SoMo phenomenon, and even the more intricate ones, like the chemotactic index, do not allow the quantification of the complete migratory behavior of the system.

Therefore, we propose a quantification method solely based on colony morphology, independent of other measured variables, and applicable across all demonstrated SoMo assays. This method builds on metrics introduced for yeast colonies [10] and assigns numerical values to colonies, indicating their degree of SoMo activity, rather than relying on categorical statements. To get a better understanding of the measurements, we introduced simulated colonies based on a modified version of the Eden model [11]. This combination of quantification and simulation of colony morphology enhances the comparability between experiments and allows for a more nuanced exploration of the influence of different parameters on SoMo dynamics. Our pipeline is scale-free, utilizing segmented image data and is openly available on GitHub and easily adaptable by other research groups.

## 2. Methods

We developed an image analysis pipeline to quantify the changes in colony morphology observed in microscopy images (Fig.1). To facilitate comprehension of the findings, we generated artificially grown colonies based on the Eden growth model [11] representing extreme scenarios across various growth regimes and compared their behavior with that of real colonies using adapted metrics initially introduced by Binder et al. [10] for the quantification of spatial growth patterns of yeast colonies.

**Figure 1.**
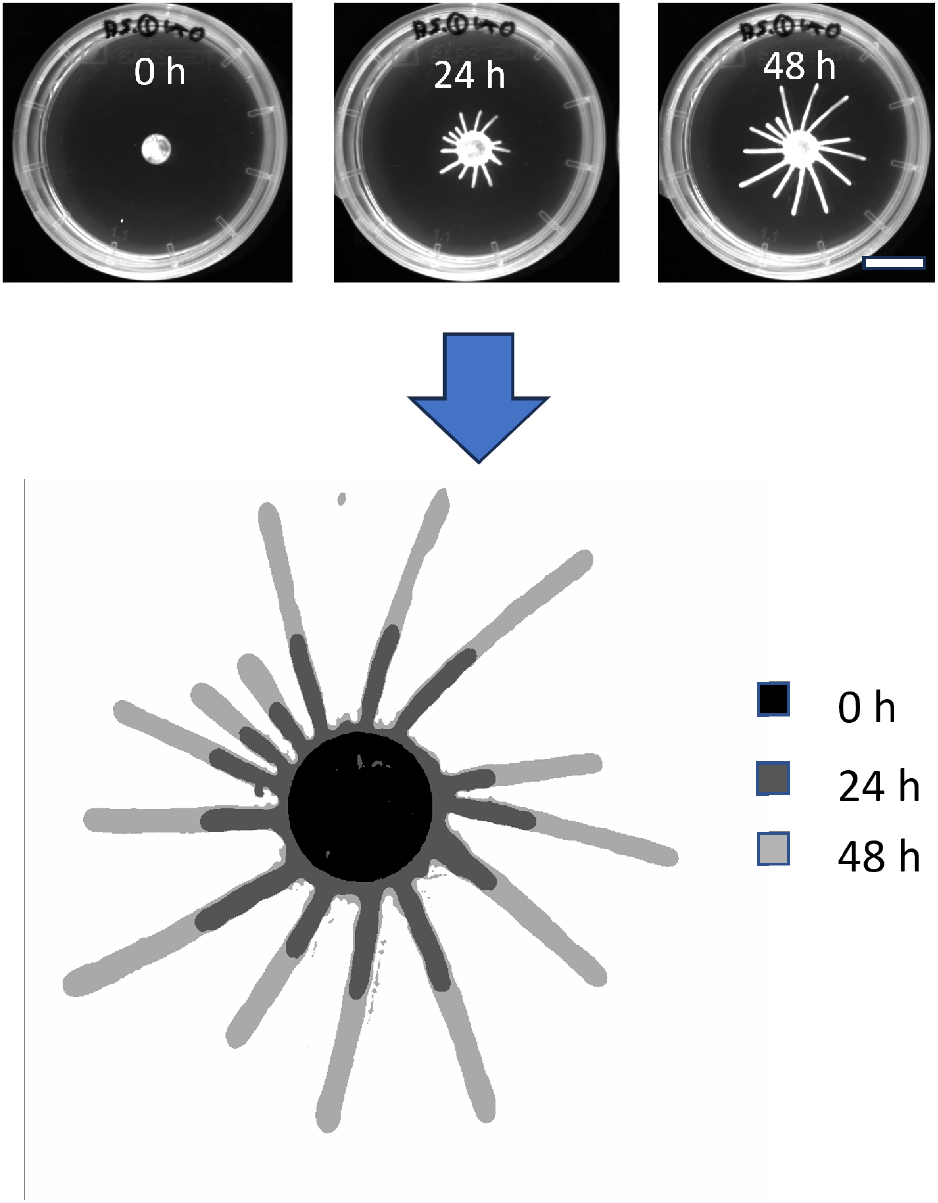
Image analysis pipeline transforms time series fluorescence images (top) into segmented and aligned stacks of binary images (bottom) for further processing. Time points are indicated by pseudocoloring. scale bar 20 mm

**Figure 2.**
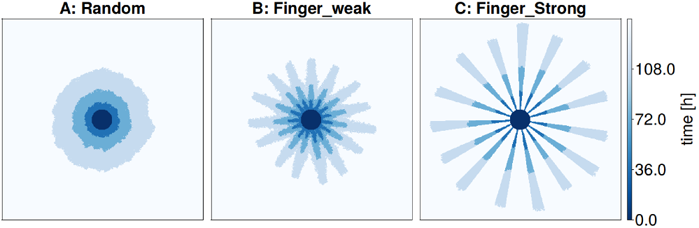
Examples of artificially created colonies. A) shows the classical Eden model with solely random growth, B) is a mixture of random growth and growth restricted to 15 directions and C) allows growth only in 15 restricted circular sections. All models are normalised to the same amount of area growth as obtained from the previous section.

### 2.1 Cell culture, motility assay and image acquisition

Cell culture and motility assays were performed essentially as in [9] with the following specifics. AnTat 1.1 and AnTat 1.1 tdTomato cells were grown in SDM-79-glc supplemented with 10% fetal bovine serum (FBS). 20% of fluorescent cells (expressing tdTomato) were added to the wildtype AnTat 1.1 cells before each experiment. The trypanosomes were concentrated to a density of 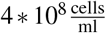 of which 2 l per colony were dropped onto an agarose gel. This concentration was calculated to result in 106 cells spread in a monolayer of 30 mm^2^ surface area, concentrated at 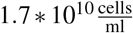 in a colony drop volume of 0.06 l after absorption of the culture medium by the hydrogel. For the experiment in Fig. 11, the initial concentration was five times the calculated monolayer amount and double the total amount of cells, resulting in faster initiation of projections. For gel production, 1 ml SDM-79-glc containing 0.4% agarose was poured in 35 mm petri dishes, left to gel 1 h with a closed lid and dry for 30 min without lid under a laminar flow hood. Gels were recorded with the iBright CL1000 imager (Thermo Fisher Scientific).

### 2.2 Image analysis pipeline

Our primary objective in quantifying colony morphology change/growth is to analyze the alteration in occupied area over time. Hence, the image analysis pipeline aims to accurately segment colonies from the background and align individual colonies across the time series (Fig. 1).

Segmentation is performed using the machine learning-based Fiji plugin, Trainable Weka Segmentation [12, 13] resulting in binary images where pixels with the value one belong to the colony and those with value zero do not. Subsequently, alignment and stacking are executed using the Rigid Body transformation type of the Fiji plugin StackReg [14]. As alignment failure occurs in approximately 20% of images, a robust yet imprecise backup approach was implemented in the programming language Julia [15]. This approach utilises convolutions to approximate the original center of the colony at *t* = 0 for all measured time-series images of a colony. For a detailed description of the image analysis pipeline, see Appendix A.

### 2.3 Artificial colonies

One of our aims was to relate the measured data to fundamental processes. Therefore, we generated artificial colonies employing a modified version of the Eden growth model[11], to enable classification of the *in vitro* data.

#### 2.3.1 Eden model

An increase in area for quasi two-dimensional colonies can occur with different characteristics. We used the area growth dynamics obtained in section3 and simulated three versions of the on lattice Eden model. The initial version (version A) adheres to the classical Eden model, featuring a completely random selection of growth sites that are empty and neighboring at least one site that is part of the colony. The second variant (version B) combines elements of the classical Eden model with partially directional growth. For a given average number of fingers, a corresponding number of circular sections with a similar width are designated as preferred growth directions with higher probability of site selection. Version C of the model builds upon this concept by restricting growth exclusively to these preferred directions. At the onset of each simulation, the directions of the circular sections are randomly generated and roughly evenly distributed across the colony surface.

Since time resolution in these artificial colonies is only limited by computation time, we chose a much more fine grained time resolution of two hour increments as compared to the experimental data. We performed simulations for 128 colonies for each version. These simulations resulted in binary images that can be directly compared to the experimental images processed by the image analysis pipeline (Fig 1). Details of the implementation can be found in Appendix C

### 2.4 Metrics

A binary image can be conceptualised as a matrix *M*(*x, y*) with

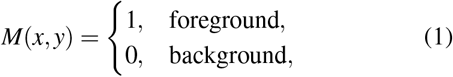

where (*x, y*) represents the position of a pixel. Additionally, we define the set of vectors *V*_*all*_, which encompasses all vectors pointing towards a foreground pixel. With these definitions, the centroid *s* of each colony can be computed as

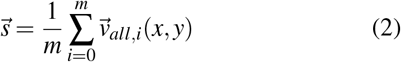

Applying these definitions to the segmented binary images, we computed binary net growth images *I*_*net*_ (*t*) that only consist of the additional area grown in a colony and their respective set of vectors *V*_*net*_. If not specifically mentioned otherwise all vectors used in further calculations are always from the set *V*_net_.

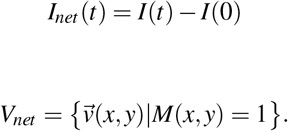

#### 2.4.1 Angular metric

The angular metric *S*_*θ*_(*i*) describes the radial variations of a colony. It is calculated by dividing the net growth binary images *I*_*net*_ into *N* circular sectors (Fig. 3) and counting the number of occupied pixels inside each sector (Fig. 4). The centroid 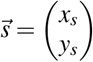 of the initial image of a colony serves as the center of the circular sectors for all images in one time series. We get

**Figure 3.**
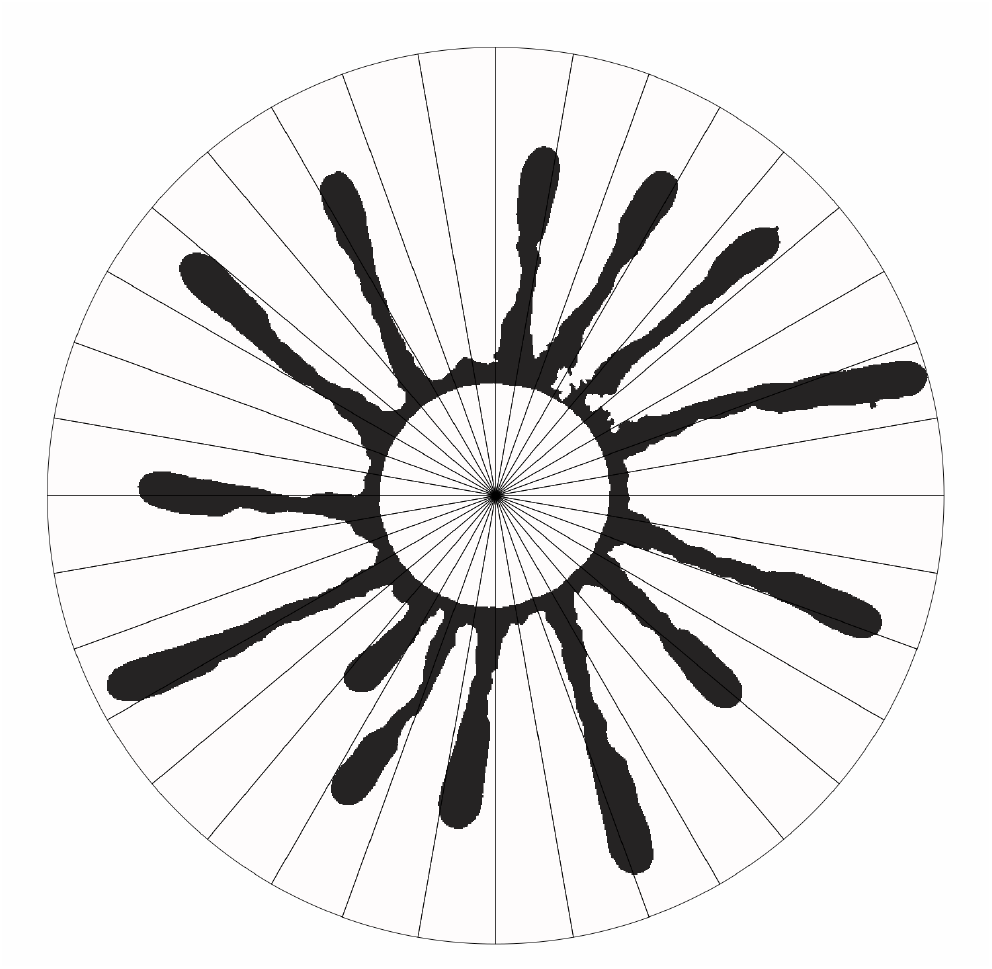
Example net growth image *I*_*net*_(48*h*) of a colony divided into 36 circular sectors (for illustration) originating from the center of the original circular shaped colony at *T* = 0*h*.

**Figure 4.**
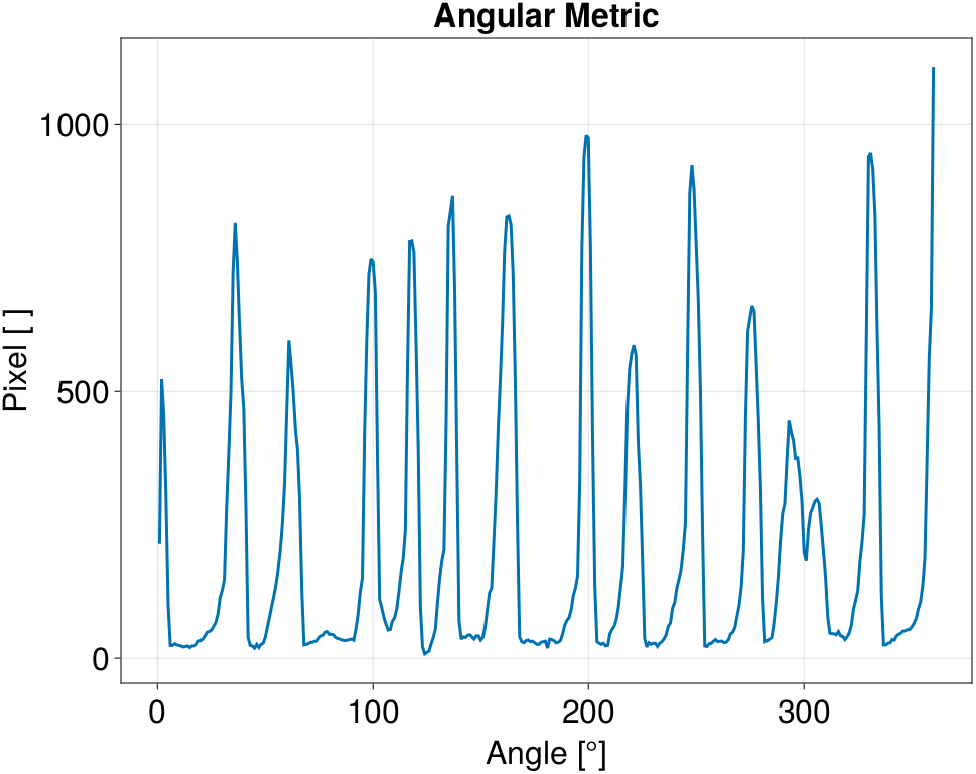
Angular metric for N = 360, for the net growth area of the colony shown in Fig.3. Each of the peaks corresponds to a finger in the image.

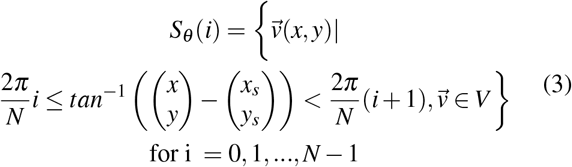

In this work, we chose *N* = 360.

#### 2.4.2 Pair correlation metric

The pair correlation metric *S*_*σ*_ (*i*) describes the angular density variations in the colony based on the net growth image (Fig. 5). All possible pairwise combinations of vectors to foreground pixels are generated and then binned with respect to the angle between them. We get

**Figure 5.**
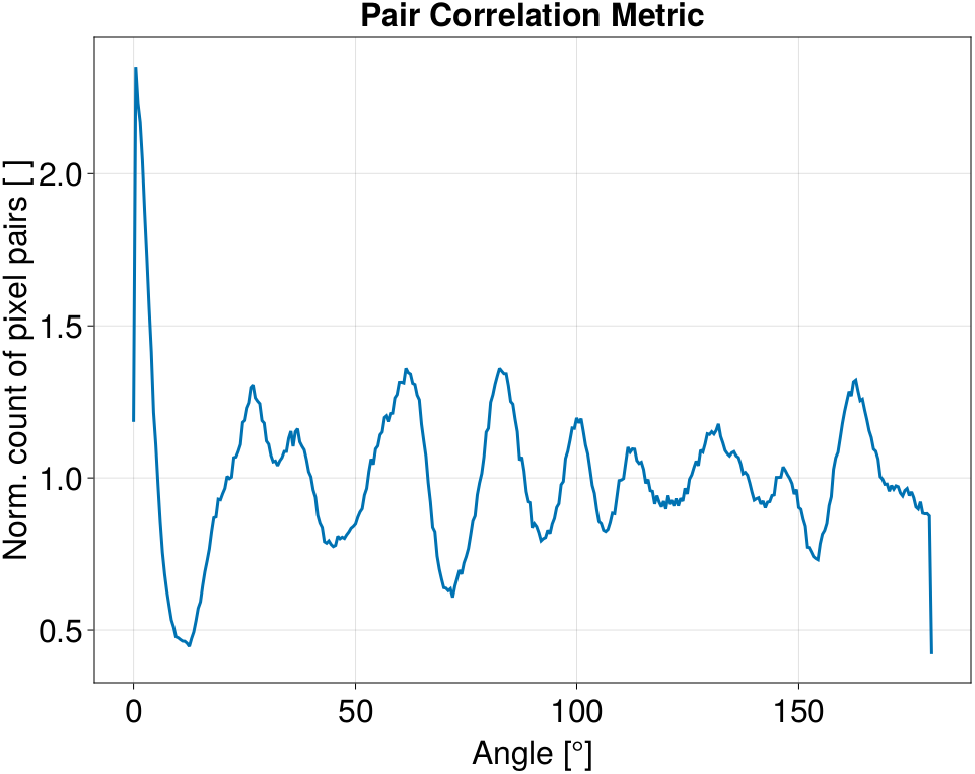
Pair correlation metric for *N*_*p*_ = 180, for the net growth area of the colony shown in Fig. 3.

**Figure 6.**
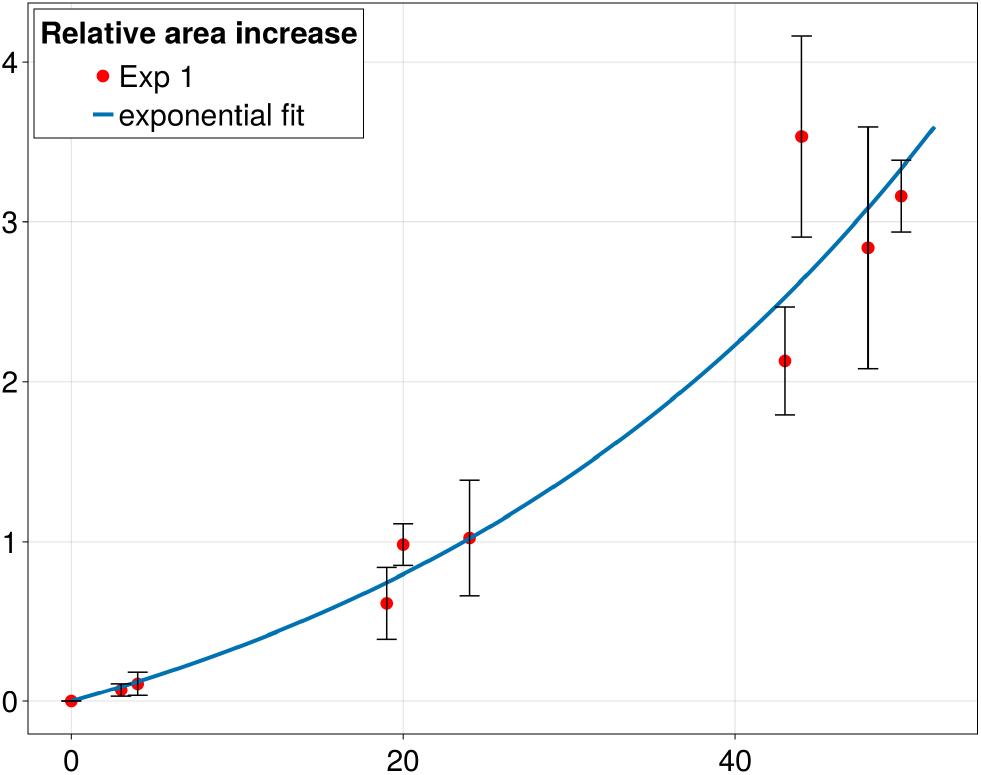
Mean relative area increase over 48 h in 32 colonies from data set Exp 1 is fitted with an exponential. The error bars represent the standard deviation.

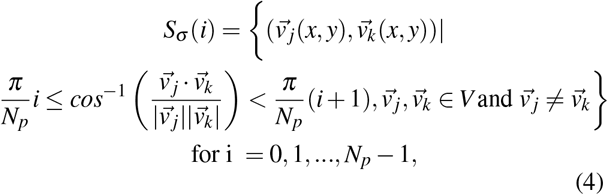

where *N*_*p*_ is the number of angular sectors. In this work, we chose *N*_*p*_ = 180.

To enable comparison of the pair correlation metric between images of different sizes, the count of vector pairs in each sector is normalised by the total number of vector pairs 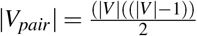 divided by the number of angular sectors *N*_*p*_. Given that |*V*_*pair*_| scales with *O*(|*V*|^2^), computational efficiency becomes increasingly challenging for larger images. Consequently, an expedient strategy is employed, wherein only a subset of randomly selected vector pairs, denoted as *V*_*sub*_, is evaluated [10]. We have empirically determined that if

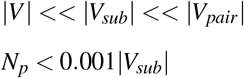

it is

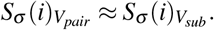

This reduction to a subset, *V*_*sub*_, allows for a practical approximation of the pair correlation metric *S*_*σ*_ (*i*) while ensuring computational tractability, especially for large images.

The distinctive peaks observed in the graph (Fig. 5) denote the normalised count of pixel pairs for which the vectors align at the specified angle. For a perfect spherical distribution of pixels, this frequency would be constant across all angles. However, in the context of a finger-like geometry, the primary and most pronounced peak signifies the accumulation of pixel pairs along the same finger. Subsequent peaks, with interspersed distances, represent the average separation between a finger and its nearest, next-nearest, and subsequent neighbors.

The height of the first peak in the normalized pair correlation metric (Fig. 5), indicates an elevated probability of locating another pixel vector at the same angle compared to a random angle. Since in our context, the first peak is always the largest, we used

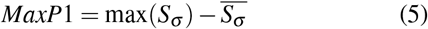

for a quantitative characterisation. Hence, *MaxP*1 is the relative surplus of the maximum over the mean of the pair correlation metric.

### 2.5 Implementation

Unless otherwise stated, analyses and simulations were performed in Julia [15] and its package Dataframes.jl [16]. All visualisations have been created with the package Makie.jl [17]. The documented code is available at Github.

## 3. Results

### *In vitro Trypanosoma* colonies show formation of finger-like structures

We placed droplets of trypanosomes in suspension onto agarose plates and imaged their evolution over 48 h (Fig 1). The colonies show a clear formation of finger-like structures. As a basis for the subsequent quantitative analyses, we performed segmentation and registration of the colony images. We performed four experiments with eight colonies each, resulting in a data set consisting of 32 colonies that were imaged at three to six different time points. This data set used in all following analyses and labelled Exp 1.

### *Trypanosoma* colonies exhibit an exponential increase in area

First, we analysed the relative area increase of the *Trypanosoma* colonies over time 6. The measured area was normalised in all colonies to the area of each colony at *A*(*t* = 0) = *A*_0_. Given the typical monolayer arrangement of *Trypanosoma* colonies in our experiments has a consistent density, it is reasonable to assume that the increase in colony area *A* is directly proportional to the increase in the count of cells. Assuming that nutrient concentration is not a limiting factor within the observed time spans, we presume exponential growth in the number of trypanosomes with growth rate *r*, hence

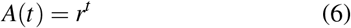

The exponential fit results in a doubling time of *θ* = 1/*r ≈* 33.65 h for the area of the *Trypanosoma* colonies.

### Quantitative analyses suggest dynamic changes in the expansion behavior of *Trypanosoma* colonies

The pair correlation metric describes the angular density variations in the outer region of the colonies. We applied it to the experimental as well as artificial data generated by a variation of the Eden model and calculated the relative maximum peak height *MaxP*1 (Fig. 7). The Eden model describes the expansion of clusters from predetermined seeds through the stochastic accumulation of particles on their surface. While originally formulated to simulate the growth of bacterial colonies [18, 19, 20], the Eden model has found applications in diverse biological systems, encompassing processes like tissue accumulation in wound healing[21]. Moreover, it extends its utility to the growth of clusters in pure physical systems, such as the formation of charged particles[22].

**Figure 7.**
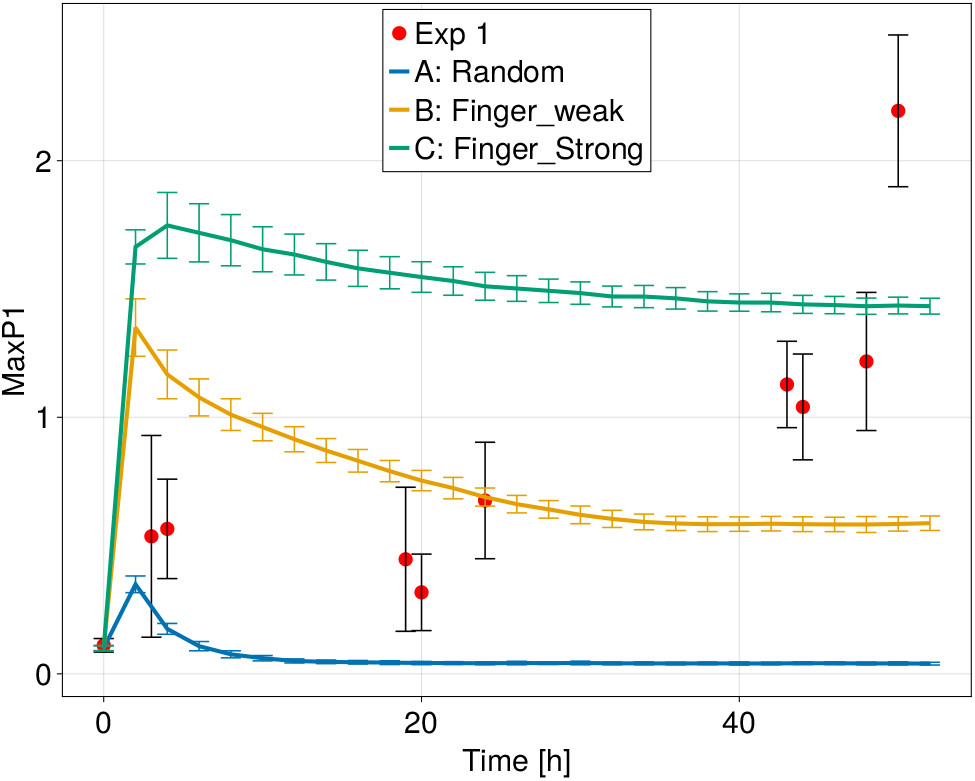
The relative maximum peak height *MaxP*1 of the pair correlation metric for the three versions of the Eden model and the experimental *in vitro* data. The data points are shown as mean *±* standard deviation. The lines are connecting the data points.

**Figure 8.**
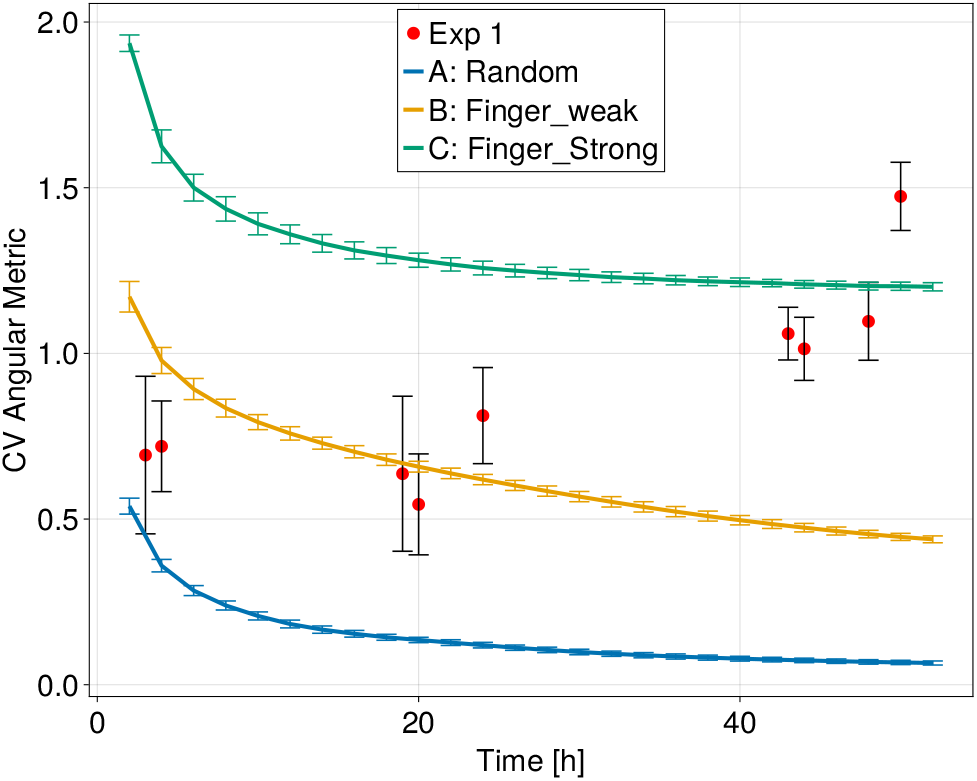
The coefficient of variation *CV* of the angular metric for the three versions of the Eden model and the experimental *in vitro* data. The data points are shown as mean *±* standard deviation. The lines are connecting the data

We find for the artificially created colonies that after an initial transition phase, the value for *MaxP*1 approaches a constant value. For version A of the Eden model, *MaxP*1 approaches a value close to zero, as the pixels are very close to a spherical distribution without an elevated probability to find more pixels under a specific angle. In contrast, version B, random growth in combination with directed fingering, displays *MaxP*1 values ranging between 0.8 and 0.5. Notably, version C, the directed finger growth, maintains the highest *MaxP*1 values exceeding one consistently. The initial peak in all three version can be explained by the limited amount of added pixels in the first time steps, which naturally creates a small anisotrophy in their distribution.

The *in vitro* colonies show a very different behavior where *MaxP*1 exhibits an increase over time, in contrast to the static values observed in *in silico* created colonies. This suggests a dynamic change in the expansion behavior of real systems. Interestingly, the value of *MaxP*1 for real colonies, while similar to the hybrid Eden model B for small time points, gradually increases and first approaches and then exeeds the values observed for version C of the Eden model with restricted growth directions.

To test these results, we next performed analyses based on the angular metric. It describes the radial variation of the colonies. As such it can be interpreted as the periodic finger growth signal. Then, perfect circular growth corresponds to a constant signal and more finger-like growth creates increased fluctuations. A standardised measure of dispersion in signal analysis is the coefficient of variation (*CV*) that is defined as the ratio of the standard deviation over the mean 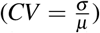 [23]. In the context of the trypanosome colonies, the *CV* becomes a quantitative measure for evaluating the “fingerlikeness” of a colony.

We calculated the *CV* for both the *in vitro* colonies as well as the artificially generated data (Fig 10). Similar to the *MaxP*1, the *CV* shows an asymptotical decrease for all three version of artificially created colonies (Fig. 10). This is again due to the random growth together with the low number of added pixels for smaller time steps, which initially creates a larger anisotropy in the distribution of growth sites.

Version A of the Eden model approaches zero for bigger time steps, which is expected as the signal becomes more and more isotropic for larger pixel numbers. Version B assumes larger values between 1.2 and 0.4 as the partially restricted growth prevents the signal from becoming constant. Version C assumes the highest values between 1.9 and 1.2, as the pure restricted growth has the largest fluctuations in the angular metric.

The real colonies show a different behavior, where the *CV* initially assumes similar values as for version B but increases over time and assumes values larger than for version C.

Considering the angular metric as the periodic finger signal suggests a Fourier transformation for further analysis. This produces the spectral density 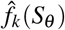 of the finger growth signal. The normalised absolute value/amplitude 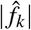 of the individual frequencies is calculated. The amplitude is normalised by the area of each individual colony at *A*(*t* = 0) = *A*_0_. Hence, we get

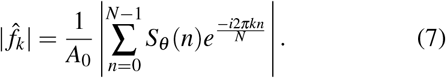

As the first component of the amplitude corresponds to the offset of the signal from zero which is proportional to the net grown area of the colony, it is neglected in all further analyses. In line with that, the mean of all amplitudes *I*_*k*_ for *k* > 1 is calculated as

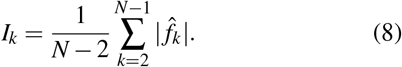

*I*_*k*_ can be interpreted as a measure for the amount of fluctuations of the surface of a colony.

*I*_*k*_ only increases slightly over time in the Eden model in version A, which grows almost in a constant uniform circular fashion (Fig. 9). Version B shows a much stronger increase over time and version C the strongest as it exclusively grows in a non-circular way. The *in vitro* colonies show again a dynamic transition between the artificial cases. Initially their *I*_*k*_ starts near 0 as version A and B, after about 20 hours *I*_*k*_ is between B and C and after about 48 hours *I*_*k*_ exceeds version C.

**Figure 9.**
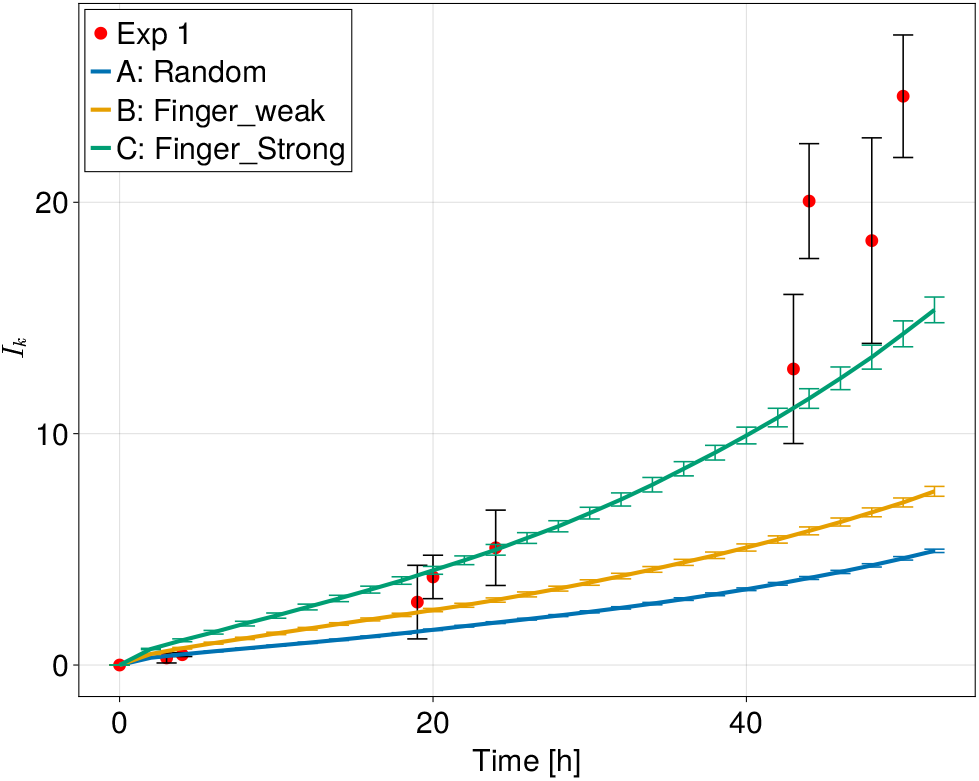
The normalised average amplitude *I*_*k*_ of the Fourier transformed angular metric for the three versions of the Eden model and the experimental *in vitro* data. The data points are shown as mean *±* standard deviation. The lines are connecting the data points.

**Figure 10.**
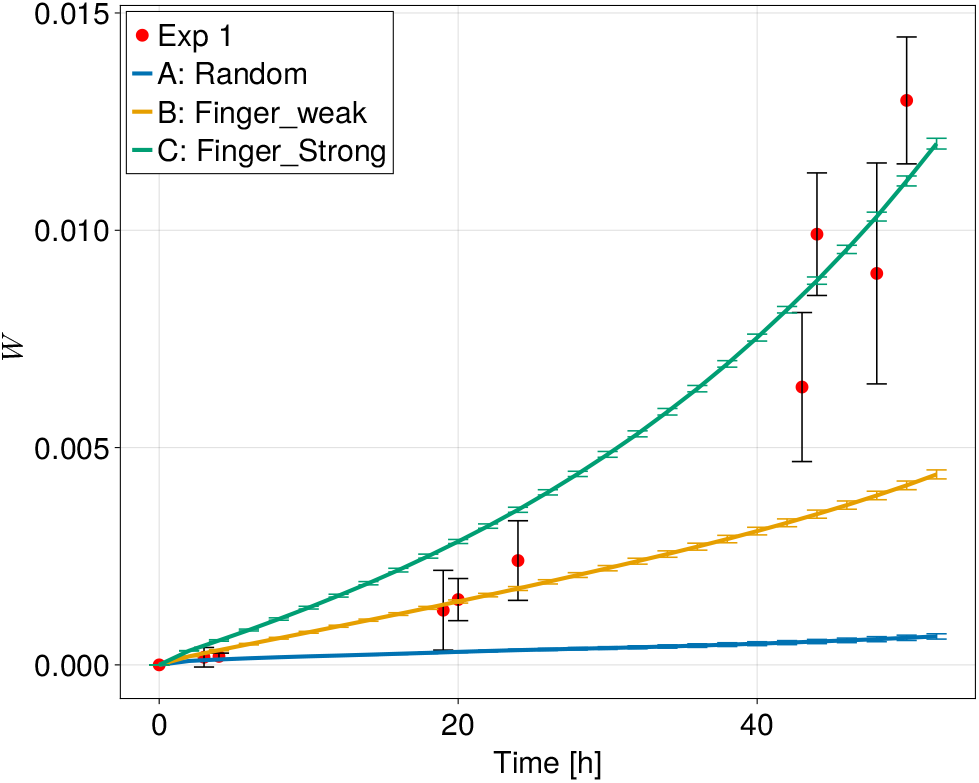
The surface roughness for the three versions of the Eden model and the experimental *in vitro data*. The data points are shown as mean ± standard deviation. The lines are connecting the data points.

**Figure 11.**
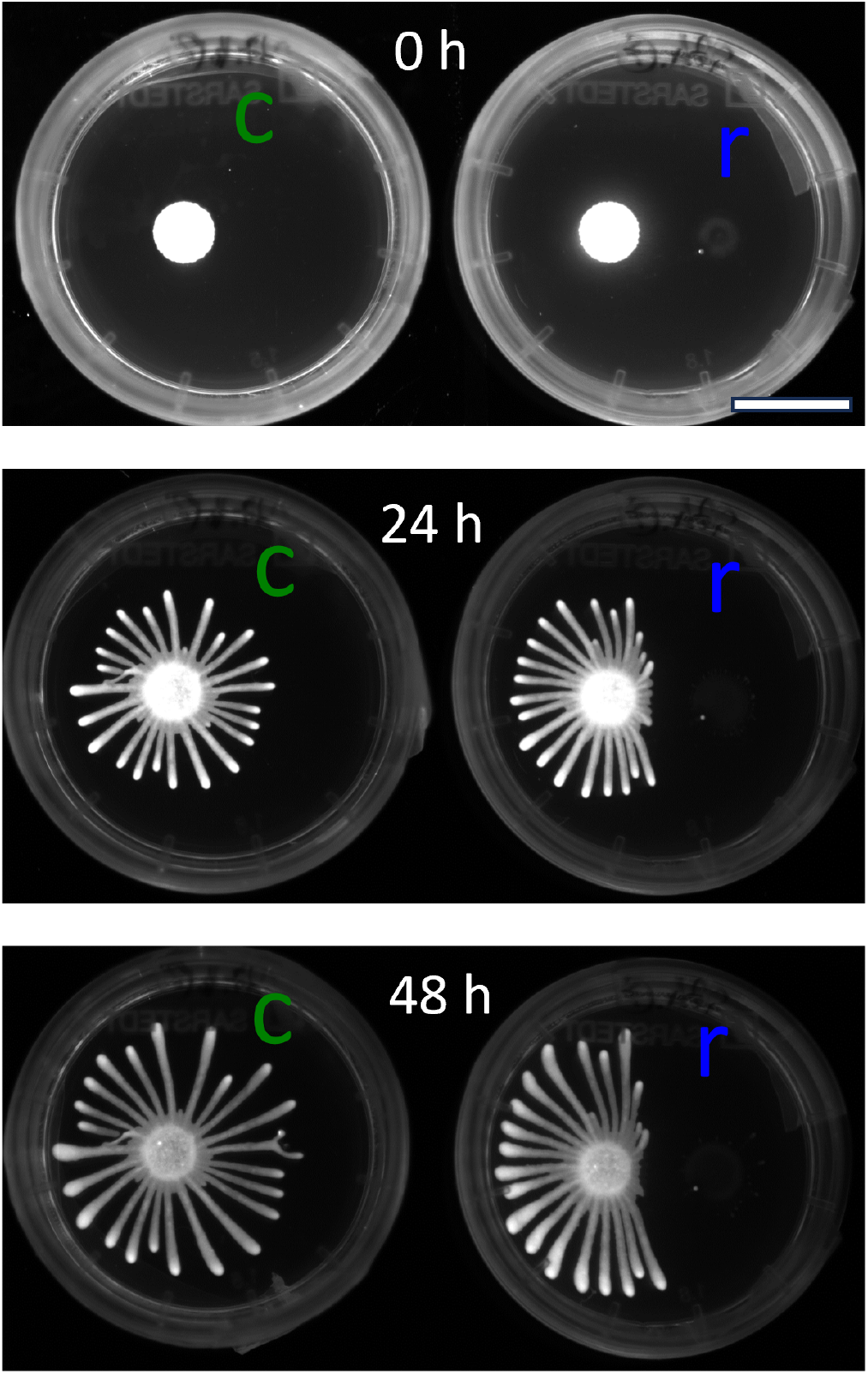
Time series fluorescence images of two *Trypanosoma* colonies from a single assay (Exp 2). The colony on the right (r) is hindered in its expansion by a SoMo negative colony, while the left control colony (c) expands regularly without hindrance. Scale bar: 20 mm

A second possible interpretation of the angular metric is as the grown surface of the colony projected into one spatial dimension. Hence, metrics to quantify surfaces can be applied. A typical quantity to characterise surface growth processes/roughness is the root-mean-squared deviation of the height *h*(*x*), often called the global interface width *W* [24]. To make this surface roughness independent of system size, we normalized the height measurements by the size of the area of each individual colony at *A*(*t* = 0) = *A*_0_ and obtained

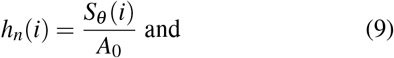

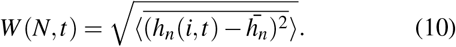

The overbar denotes averaging over all i in a single system of size *N* and the ⟨-⟩denotes averaging over an ensemble of *n* systems.

The Eden model in version A shows only a small increase in *W* in the observed time frame. The versions B and C are far away from this regime and their roughness shows a stronger increase over time. This increase in roughness is stronger in version C. The *in vitro* colonies show again a dynamic switch, but this time with a closer resemblance to version C of the Eden model for late time points. Hence, their roughness is initially close to 0, then after about 20 hours most similar to version B, and after about 48 hours most similar to version C of the Eden model.

In summary, all four measurements show a clear distinction for the three different versions of the Eden model. Comparing these results to the measurements for the *in vitro* data suggests that the *Trypanosoma* colonies first grow randomly and then restricted growth within specific directions becomes increasingly dominant.

### Growth phases are robust with respect to number of cells and inhibition

To assess the robustness of different growth phases, an additional assay was conducted involving six colonies. Two colonies were subjected to repulsion by a SoMo negative trypanosome colony (Exp 2: repelled), while four served as control colonies without repulsive signal (Exp 2: control).In addition, for all six colonies the initial cell number was larger than in the previous experiments. This led to accelerated expansion and the formation of a larger number of fingers (approximately 30) (Fig.11).

These colonies exhibited slightly different area growth dynamics due to their initial non-monolayer configuration. We employed a modified exponential growth model to fit the area data and calculated the relative area increase over time (see Appendix B for further details).

We simulated colonies with these modified growth dynamics and 30 fingers, then applied our established metrics to the additional data. We find that *W* and *I*_*k*_ exhibited an increase for colonies with more fingers, while *MaxP*1 and *CV* decreased (Fig. 12). This trend held true for both simulated and real colonies. The peaks in these measurements just before 20 h relate to the altered growth dynamics of the colonies with more cells (see Appendix B for further details). We further find that control colonies display a transition behavior, becoming more similar to version B of the Eden model (with 30 fingers) after 24 hours and more like version C after 48 hours. A slight difference between control and repelled colonies was observed, albeit inconclusive. Hence, an alternative quantification of this geometric change is required.

**Figure 12.**
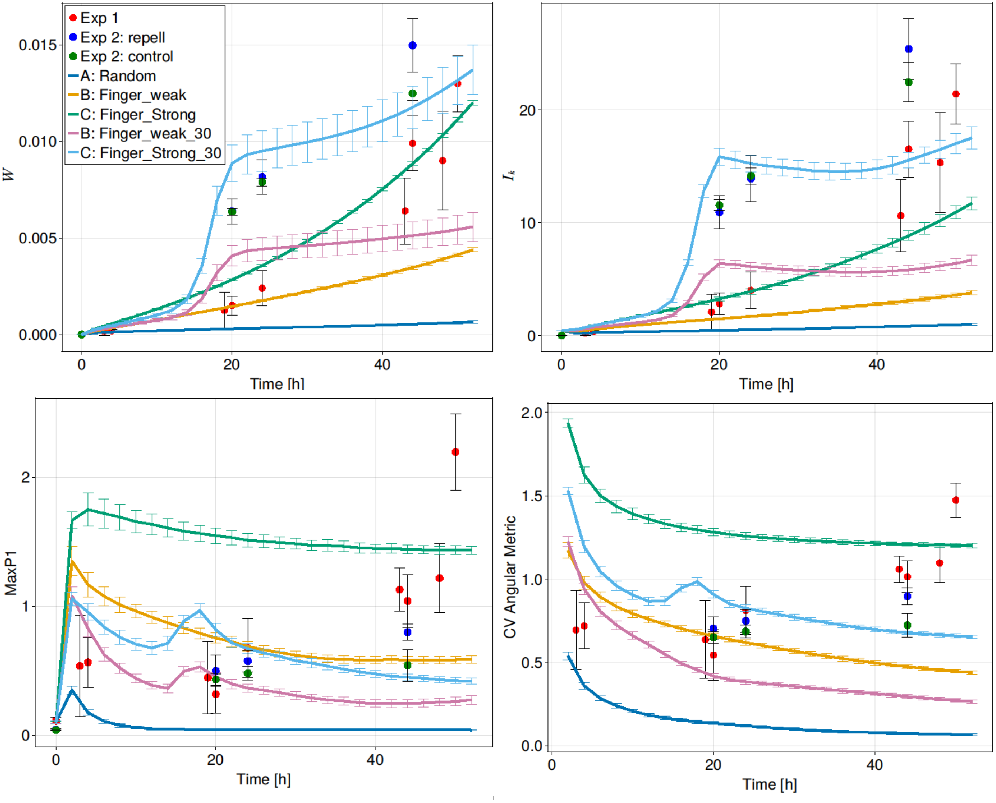
*W, I*_*k*_, *MaxP*1,*CV* are shown for the previous simulated and *in vitro* colonies (Exp 1) together with control, repelled (Exp 2) and simulated colonies with 30 fingers and accelerated expansion.

### Quantification of deflection

To quantify the deflection, we examined the pair correlation metric *S*_*σ*_, which assesses the anisotropy in the pixel distribution (Fig. 13a). We find a distinct differentiation between control and repelled colonies, characterized by an upward shift for small angles and a downward shift for larger angles. To pinpoint this further we smoothed *S*_*σ*_ with a moving average (MA) of *S*_*σ*_ for a window of 90° (Fig. 13 b)).

**Figure 13.**
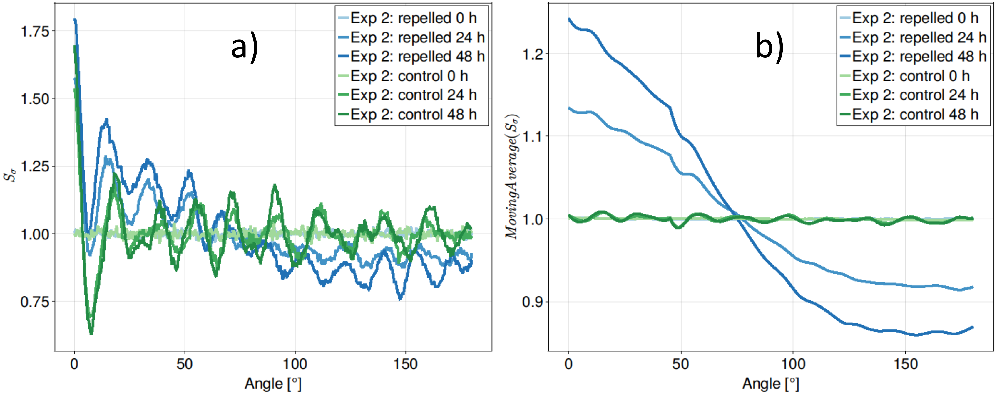
a) The mean pair correlation metric *S*_*σ*_ of repelled and control colonies of Exp 2. b) The moving average of the mean pair correlation metric *S*_*σ*_ for a window of 90° (MA) shows a distinct difference between repelled and controlled colonies.

**Figure 14.**
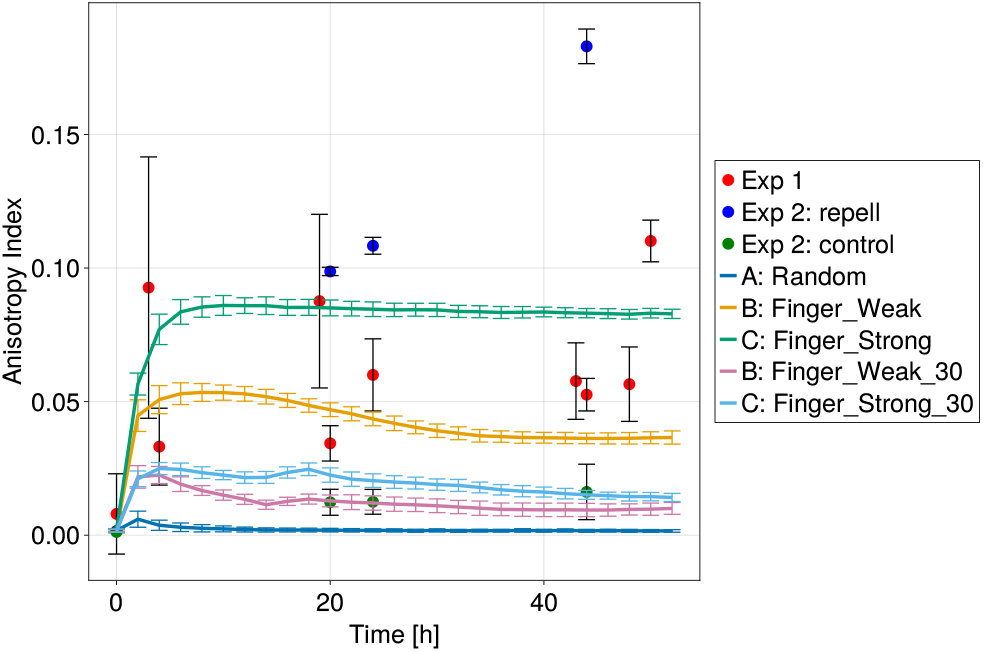
The anisotrophy index for the three versions of the Eden model, with 15 fingers and with 30 fingers, respectively, as well as the experimental *in vitro* data. The data points are shown as mean *±* standard deviation. The lines are connecting the data points.

We propose the anisotropy index *A*_*I*_, quantifying the relative relationship between the maximum and minimum values of the smoothed pair correlation metric *S*_*σ*_ as

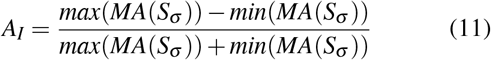

We computed the mean *A*_*I*_ for all repelled and control colonies in the additional assay, alongside the four previous assays and the simulated colonies. Given the distinct expansion behavior observed in the additional assays (faster with more fingers), we simulated two additional versions of the Eden model (B: Finger weak 30 and C:Finger Strong 30) to account for these properties.

We note that the anisotropy index *A*_*I*_ tends to be larger for colonies with fewer fingers (both real and simulated) (Fig.14), which is reasonable considering that fewer fingers indicate a less isotropic distribution of pixels. The real colonies exhibit values ranging between 0.05 and 0.1, akin to the simulated colonies of models B and C. For the last time point, the real colonies have a surprisingly large anisotropy value. This result matches with visual inspection of the corresponding images. To investigate whether this is an artefact or a relevant behavior requires more experimental data for this specific time point. Conversely, for colonies with more fingers, *A*_*I*_ is smaller, with control colonies displaying similar values to the simulated ones with 30 fingers, hovering around 0.01. Notably, repelled colonies exhibit the highest values, surpassing 0.1 after 24 hours and reaching 0.16 after 48 hours, clearly distinguishing them from the control colonies.

## 4. Discussion

We established quantification methods for *Trypanosoma* social motility assays. Our measurements allow objective evaluation of colony and finger growth dynamics as well as anisotropies. We find that the trypanosome colonies exhibit a dynamic growth process that transitions from initial pure circular expansion to exclusive “finger” expansion. These growth phases are independent of the initial number of cells and anisotropies in colony expansion.

We analysed the change in morphology of SoMo exhibiting trypansoma colonies on agarose. Fluorescence microscopy images were processed in a multistage image analysis pipeline to enable calculation of the two one-dimensional metrics angular metric and pair correlation metric introduced by Binder et al [10]. To better classify the results, we measured the area increase over time and simulated three different randomly growing colonies based on the on-lattice Eden model growth model [11]. The area doubling time that we obtained from our fitting is larger than the population doubling time reported in the literature ([8, 4]). This indicates that cell division does not double the area occupied by the cells. Instead, closer packing seems to occur. The angular and pair correlation metrics were further processed by calculating five quantities (*MaxP*1,*CV,W, I*_*k*_, *A*_*I*_) that map the growth processes to scalar values. Calculating these measures for different Eden models reveals that the measures are lowest for random growth and increase with increasing contribution of finger growth.

The different measurements exhibit a temporal evolution. For the simulated colonies, the relative maximum peak of the pair correlation metric *MaxP*1 and the coefficient of variation of the angular metric *CV* decrease over time. This can be explained by the constant growth rules in the simulated colonies which cause both these quantities to asymptotically approach a constant value for *t →* ∞. In the real colonies however both these quantities increase over time which is caused by the changing expansion behavior from spherical to finger like. Both *MaxP*1 and *CV* are sensitive to a higher number of fingers and show overall lower values due to the less pronounced single fingers compared to the average signal. Inhibition does not have a significant impact on their values.

The surface roughness *W* and normalised Fourier coefficients *I*_*k*_ even though measuring quite different quantities of the system showed very similar qualitative behavior. Both increased over time for the simulated colonies. The transition from random circular to finger-like growth behavior in real colonies can again be mapped here. Contrary to *MaxP*1 and *CV*, both *W* and *I*_*k*_ increase with increasing numbers of fingers. Inhibition also does not have a significant on their values.

Since the previous four quantities failed to distinguish between normal and inhibited colonies, a fifth quantity was developed: the anisotropy index *A*_*I*_. This metric assigns a numerical value to the anisotropy of a colony. While it excels at differentiating between inhibited and normal colonies, it may also prove useful for comparing the uniformity of growth between colonies. This allows for the accounting of anisotropies in the experimental setup, variations in cell behavior, and other factors.

Taken together, our methods demonstrate the possibility for an objective description of the behavior typically arising in trypanosome social motility assays. The combination with the Eden model allows categorising the behavior types observed. This approach is not limited to trypanosomes but can be driectly applied to other finger forming cells like yeast (ref) or bacteria (ref). Application of our methods to *in vitro* trypanosome colonies reveal that simple growth dynamics as represented by the Eden model are not sufficient to accurately capture social motility dynamics. However, in combination with data from single cells [9], our colony-scale quantification builds a solid basis for developing more intricate models, that could incorporate factors such as cell division, characteristics of the microswimmer motion and the properties of the colony boundaries. Hence, we are one step further in understanding the mechanisms of collective trypanosome motion *in vitro*.

## Appendices

### A Detailed description of the image analysis pipeline

The fluorescence images were preprocessed with a Gaussian filter (*σ* = 1.5). This step helps prevent false detections caused by uneven illumination, particularly in thinner and less bright regions of finger structures. The segmentation was done by the machine learning Fiji [12] plugin Trainable Weka Segmentation [13]. One of the images of a colony at *t* = 48 *hours* was used for initial training. Areas of the background and colony have been manually selected to serve as basis for the training of a classifier. After the application of the classifier on multiple colonies and time points, the selected areas were manually adapted to optimize further training.

Subsequently, the Fill Holes function provided by MorphoLibJ [25] was applied to binary images to eliminate falsely classified holes fully contained within the colony. Additionally, artifacts such as the bright edge of the agar plate, incorrectly identified as part of the colony, were manually removed.

In the next stage, binary images of the same colony were stacked and aligned (Fig. 1 in the main text) using the Rigid Body transformation type of the Fiji plugin StackReg [14]. Subsequently, the first image (*t* = 0*h*) was subtracted from later images to isolate the net grown area for further analysis.

Later analysis showed, that the pair correlation metric *S*_*σ*_ for small timesteps (*t* < 10*h*) is very sensitive to small shifts/errors in the alignment process causing anisotrophies in the net growth images *I*_*net*_, even though the colony expands perfectly circular in that time frame. If this was confirmed manually, an alternative technique was used to isolate the net grown area from the original colony structure relying solely on the information contained in a single image. This approach used the following assumptions:

- All colonies start out with an almost perfect spherical shape
- Expansion is majorly in the form of asymmetrical protrusions which rapidly evolve into distinct finger-like shapes

The first step is to calculate the convolution of the binary colony image with a rectangular kernel roughly 40% the area of the original colony (Fig 15). This convolution yields an intensity image that is binarised with a threshold of 0.8 where all pixels below that threshold are set to zero and all above to one (Fig. 16 top).

**Figure 15.**
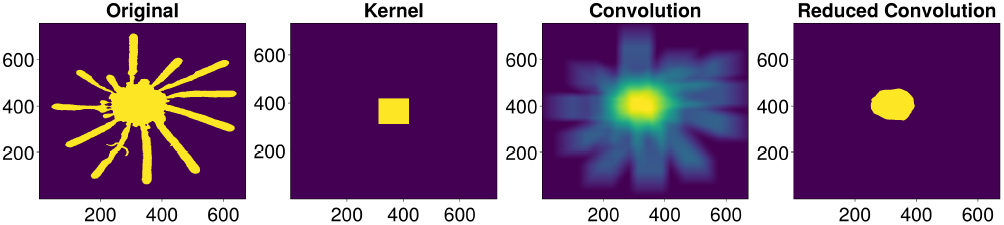
Convolution of a colony binary image with a rectangular kernel creates an intensity image which again is binarised.

**Figure 16.**
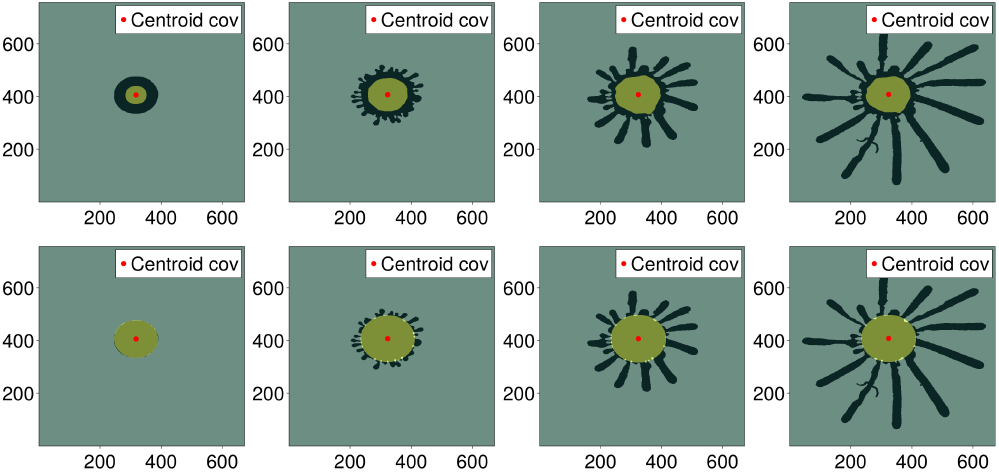
Time series of a colony shown together with the reduced colony and its centroid produced by convolution (top). and together with the fitted circle from the centroid of the reduced image (bottom).

**Figure 17.**
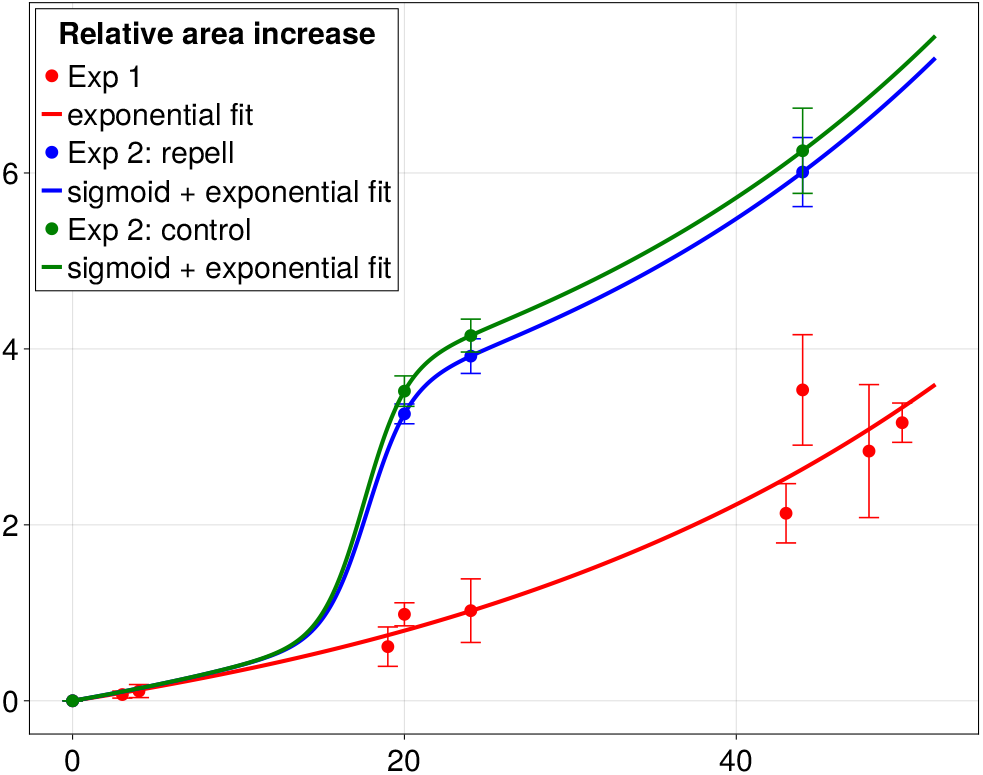
Mean relative area increase over 48 h for the three different data sets fitted with an exponential (and sigmoid functions). The error bars represent the standard deviation.

Then, the centroid 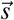 (eq.2) of the reduced convolution image is calculated and used to fit a circle into the original binary colony (Fig. 16 bottom). This circle should closely approximate the colony’s initial spherical state at *t* = 0. The circle is gradually build up from the centroid of the colony. If less then 50% of the grid points on the outer most annulus of the circle are occupied, the fitting is stopped.

This method offers an alternative approach to differentiate between the spherical “core” region and the newly developed finger-like regions. Compared to the stacking method, this approach provides less precise but more robust results. We have calculated all our results for both methods, and they did not exhibit significant differences, except for the described issues encountered during small time points in the calculation of the pair correlation metric *S*_*σ*_. Therefore, we employed the Fiji-based stacking method for all our results, where this issue did not arise, and the convolution method for the few datasets where there were alignment problems for small time steps. We meticulously reviewed all the results to ensure their validity.

### B Area fitting of colonies with 30 fingers

A sole exponential fit to model the relative area increase over time for the additional assays featuring colonies of increased cell numbers yielded unsatisfactory results. The elevated cell density in these colonies prompted a rapid initial area increase as the colonies probably thinned toward their preferred monolayer arrangement. Simultaneously, exponential growth due to cell division contributed to overall area expansion. To effectively capture this dual behavior, a composite approach combining a sigmoid-like function to represent thinning and exponential growth to model cell division was employed. This combined model demonstrated good agreement with the observed data.

The logistic function was chosen as the sigmoidal component due to its simplicity and ease of use, although this selection was somewhat arbitrary, given the limited data points available to precisely characterize thinning-induced area increase. However, this choice is not anticipated to significantly impact subsequent analyses, which predominantly focus on time points post-thinning.

The composite fitting function is expressed as:

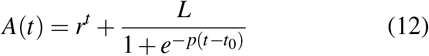

The exponential segment of the fitting function yielded doubling times of approximately *θ* = 1/*r ≈* 29.75 hours for the repelled colonies and *θ* = 1/*r ≈* 29.67 hours for the control *Trypanosoma* colonies.

### C Implementation of the Eden model

The Eden model was implemented in the programming language Julia [15], enabling growth simulations of 30 colonies with up to 10^6^ added sites in under 10 minutes. The chosen variant of the model tracks all perimeter sites, defined as empty sites with at least one surface site as a neighbor. A random selection process designates one of these sites for population, followed by an update of the perimeter sites list. Algorithmically, this operation can be executed through the convolution of a Laplacian kernel with the lattice. The result is an auxiliary lattice with the same size as the original lattice. All perimeter sites in the auxiliary lattice assume positive values corresponding to their adjacent surface sites, surface sites assume the negative count of adjacent perimeter sites, and all other sites assume a zero value (Fig 18).

**Figure 18.**
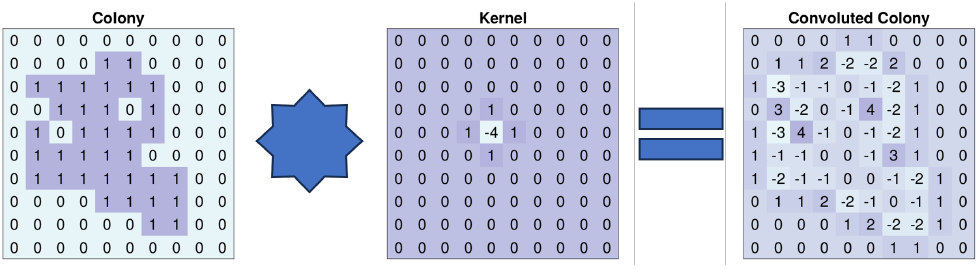
The convolution of a laplace kernel with the colony lattice extracts all perimeter and surface sites. Perimeter sites assume positive values corresponding to their adjacent surface sites. Surface sites assume negative values corresponding to their adjacent perimeter sites.

The auxiliary lattice is subsequently filtered to extract only those sites with positive values, thereby isolating the perimeter sites. Upon filling one of these sites(e.g. setting its value to one in the main lattice), a single Laplacian kernel is added to the auxiliary lattice centered around the site, triggering automatic updates for all neighboring sites accordingly (Fig 19. This results in a run time 𝒪 (*M*) that is linear with respect to the size *M* of the used lattice.

**Figure 19.**
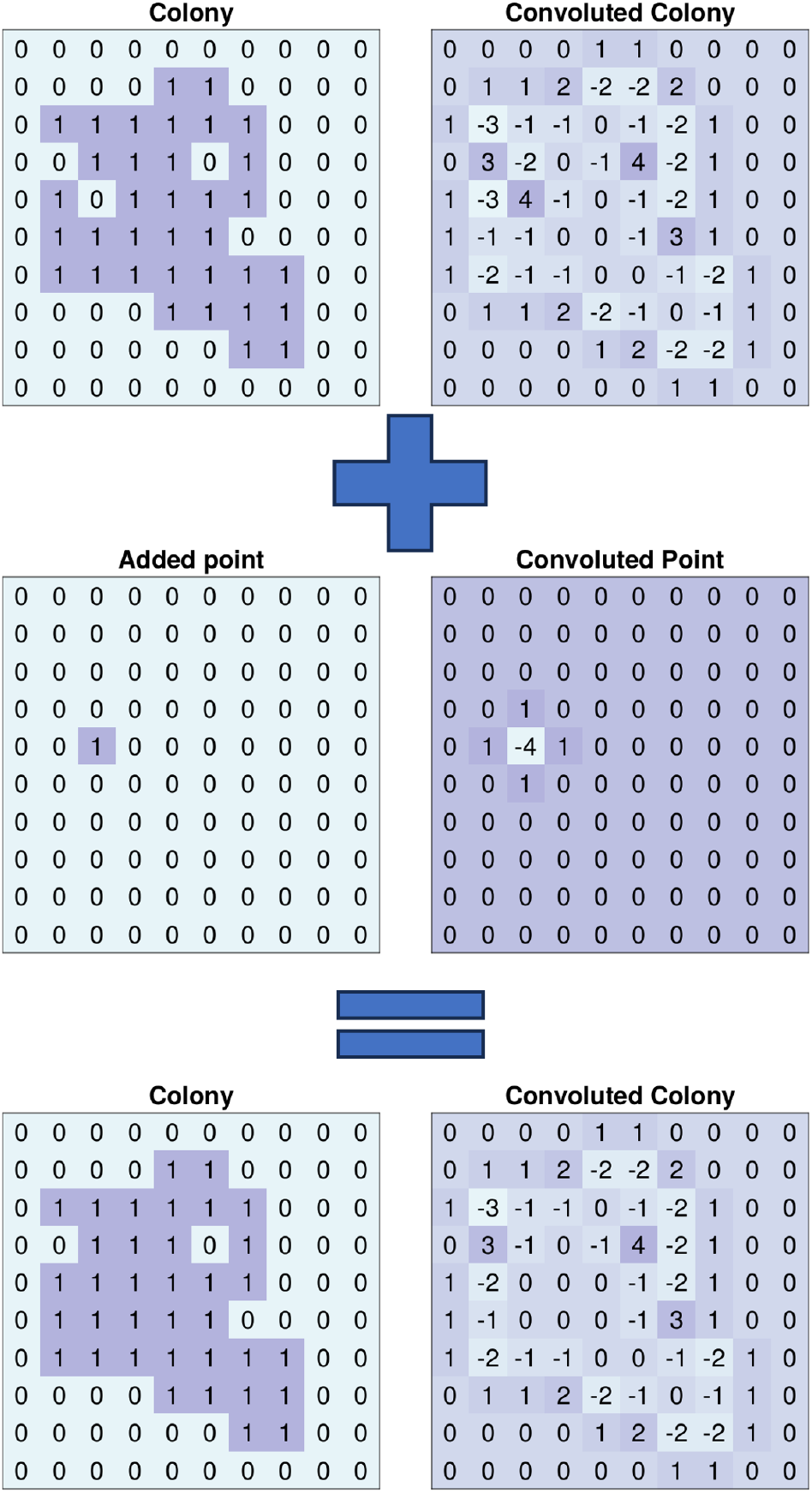
The colony and auxiliary lattice get updated simultaneously. A filled site in the colony corresponds to an added Laplace kernel at the same position in the auxiliary lattice.

